# The contributions of biological maturity and experience to fine motor development in adolescence

**DOI:** 10.1101/2024.09.02.610814

**Authors:** Andrea Berencsi, Ferenc Gombos, Lili Julia Fehér, Patrícia Gerván, Katinka Utczás, Gyöngyi Oláh, Zsófia Tróznai, Ilona Kovács

**Author notes:** Corresponding author: Andrea Berencsi, HUN-REN-ELTE-PPKE Adolescent Development Research Group Budapest, H-1088 Budapest, Mikszáth tér 1., Hungary.

## Abstract

Fine motor function develops into adulthood, but little is known about the differential effects of biological maturation and experience on speed and complex sequential performance of the hand. To determine maturity levels, ultrasonic bone age (BA) was assessed in 225 adolescents (123 females; BA range: 9.9 to 17.9 years). The role of experience was evaluated based on chronological age (CA, range: 11.1 to 16.5 years), musical instrumental experience, and handedness. Multiple linear regression modeling showed that BA is the strongest predictor of sequential motor performance, while CA influenced motor speed when no musical instrumental experience was present. When present, the amount of highly specific musical instrumental experience becomes the main predictor of sequential performance.

## INTRODUCTION

The refinement of fine motor skills in adolescence holds significance for academic achievement, the transition to adulthood, and success in the workforce. Throughout adolescence, motor coordination still undergoes development, including progress within the musculoskeletal system as well as brain regions associated with motor function ^1,2^. Complex details of fine motor development have been extensively mapped in relation to chronological age ^3–7^. Nevertheless, the interaction between biological maturation and chronological age, along with the specific influence of pubertal maturation, remains unclear. In the present study, our objective is to evaluate the respective contributions of maturation, chronological age and specific motor experience to the development of fine motor performance.

As a function of chronological age, both fine motor speed and the speed and accuracy of complex finger movements increase during the teen years ^3,4,8^. The neural background of fine motor behavior has also been investigated primarily as a function of chronological age. At the regional level, motor cortices responsible for initiating voluntary movements and prefrontal areas associated with executive control undergo synaptic pruning and thinning during adolescence, which enhances efficiency ^9,10^. Primary motor cortex has a strongly connected pattern by the age of 14 years and becomes even more connected in between 14 to 26 years ^6^. At the network level, the development of long-range connections between frontal and parietal cortices, which are associated with spatial processing and motor planning ^11,12^ extends into late-adolescence, following a local to distributed network transition ^5^. Additionally, myelination enhances the speed of motor responses, thereby facilitating the execution of fine movements ^13,14^. Myelination of the corticospinal tract that is responsible for the activation of hand muscles with a crucial role in producing speed in elaborate finger movements is steep until the age of 7-12 years ^7,15,16^ then gradually continues until adulthood ^7,16^.

Pubertal maturity, as it is related to gonadal hormone levels in childhood and adolescence, determines cortical activity during motor preparation in sensorimotor areas at the functional level ^17^. Trajectories of puberty-related cortical volumetric changes in frontal and parietal cortices show associations with pubertal tempo, suggesting an independent process from merely age-related development ^18^. White matter development in motor-related areas also shows associations with pubertal changes with great individual variability ^19^. However, while fine motor development has been precisely mapped as a function of chronological age, the interplay between biological maturation and chronological age, and the sheer contribution of pubertal maturation has not been clarified yet.

Adolescence represents a pivotal phase in human development marked by the pubertal reawakening of the hypothalamic-pituitary-gonadal (HPG) axis following an initial surge of activation postnatally. The significance of pubertal timing for the development of the brain and behavior is due to the fact that the reactivation of the HPG axis not only triggers sexual maturation, physical growth, and bone mineralization, but also coincides with profound structural modifications in the brain ^18–21^, and advancements in the cognitive ^22–24^ and socioemotional ^25,26^ domains. Nevertheless, the precise interplay between the process of pubertal maturation itself and the mere passage of time or accumulated experience on brain and behavioral development remains unclear in general, not only with respect to motor development.

The uncertainty about the contribution of maturation versus chronological age or experience is due to the general absence of a reliable assessment method for gauging different maturity levels. The widely used Tanner Scale ^27–29^ relies on subjective assessments of physical markers like breast and testicle size, rendering it susceptible to evaluator bias. Moreover, it does not consider contemporary nutritional factors or secular trends in growth ^28,30–33^. Both self- and parent-reported versions of this scale also exhibit unreliability ^27,28^. Consequently, inconsistencies in developmental progress assessments contribute to disparate terminology in the literature ^34^. To address these challenges, we propose the utilisation of ultrasonic bone age as an alternative to the Tanner scale. This method has been introduced into non-clinical developmental research very recently ^24,26,35,36^ and it seems to offer improved accuracy in the assessment of pubertal maturity. This approach aids in operationalizing the temporal dimensions of adolescence more explicitly.

### The current study

In our present cross-sectional investigation into adolescent motor development, we concentrate on bone age as a critical indicator of maturity, while also considering chronological age, hand dominance and years of instrumental musical practice as fundamental markers of experience. An earlier study has shown that instrumental musical training is an all-encompassing factor in determining fine motor performance in adolescents ^35^, therefore it seems very important to investigate its relevance independently of the impact of maturation. Incorporating musical experience as a variable in our study offers enhanced clarity regarding the influence of experience within the participant cohort that received instrumental musical training, whereas the impact of maturity is anticipated to be more discernible in the group lacking such training.

Our principal aim is to initially distinguish between the two fundamental aspects of fine motor development: cortical network connectivity and the myelination process of extended neural pathways. To accomplish this, we employ two variations of a finger tapping task ^3,8,35,37,38^: (1) a sequential finger tapping task focusing on both speed and accuracy, which is associated with long-range cortical connectivity ^11,39^, and (2) an index finger tapping task assessing maximum motor speed and its association with white matter development ^13,40^. See Figure 1. for the two variations of the finger tapping task. We additionally incorporate assessments of both the dominant and non-dominant hands of participants to examine the contrasting impact of hand utilization, employing hand preference as an indicator of experience ^41^.

**Figure 1.**
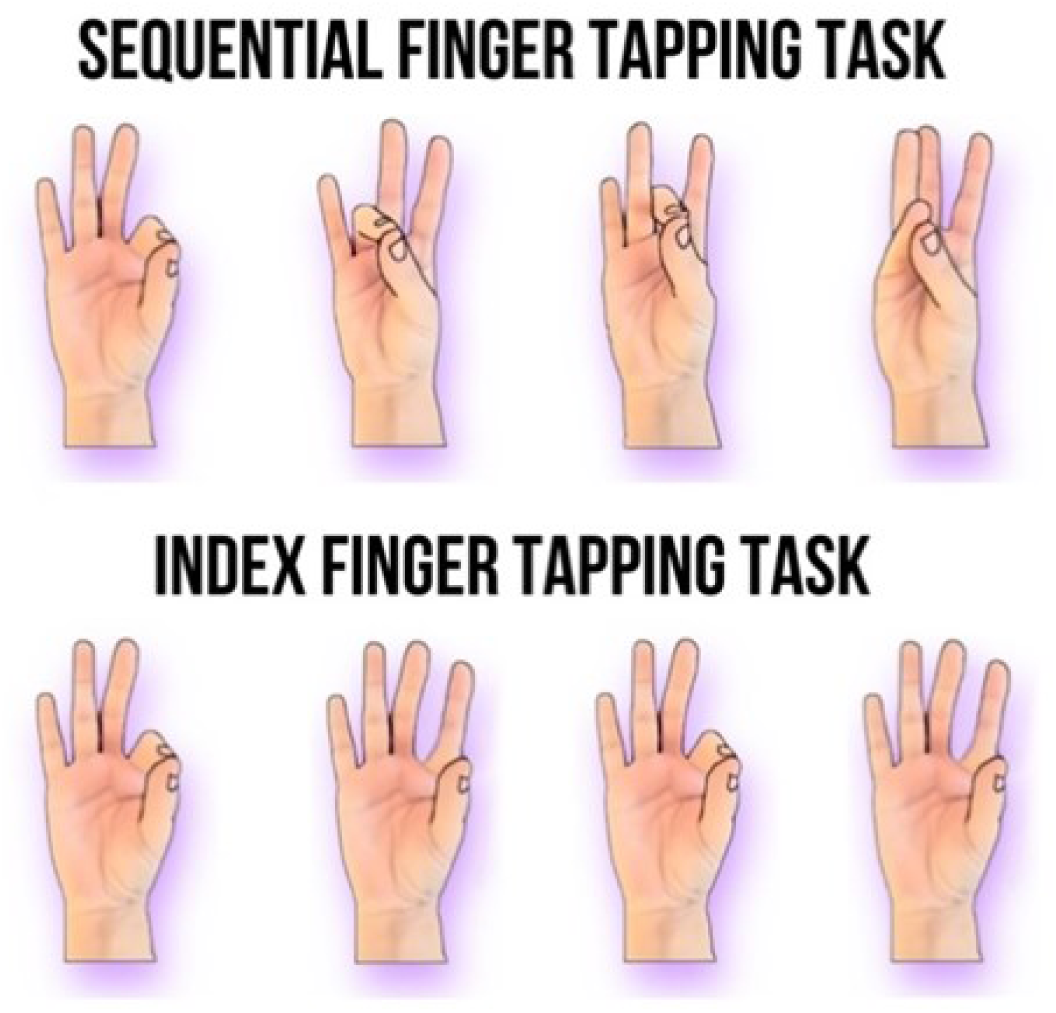
Motor development is assessed using a four-element sequential finger tapping task and a repetitive index finger tapping task, both utilizing finger-to-thumb opposition with eyes closed (refer to Methods). The sequential task focuses on executing movements as rapidly and accurately as possible, while the index finger tapping task prioritizes speed alone.

Our second aim is to disentangle the contributions of biological maturity and chronological age to the development of the above-mentioned components of motor development; therefore, we assess bone age as a proxy of individual maturation levels of the adolescent participants. As it is illustrated in Figure 2, there is a substantial variability in individual maturity levels.

**Figure 2.**
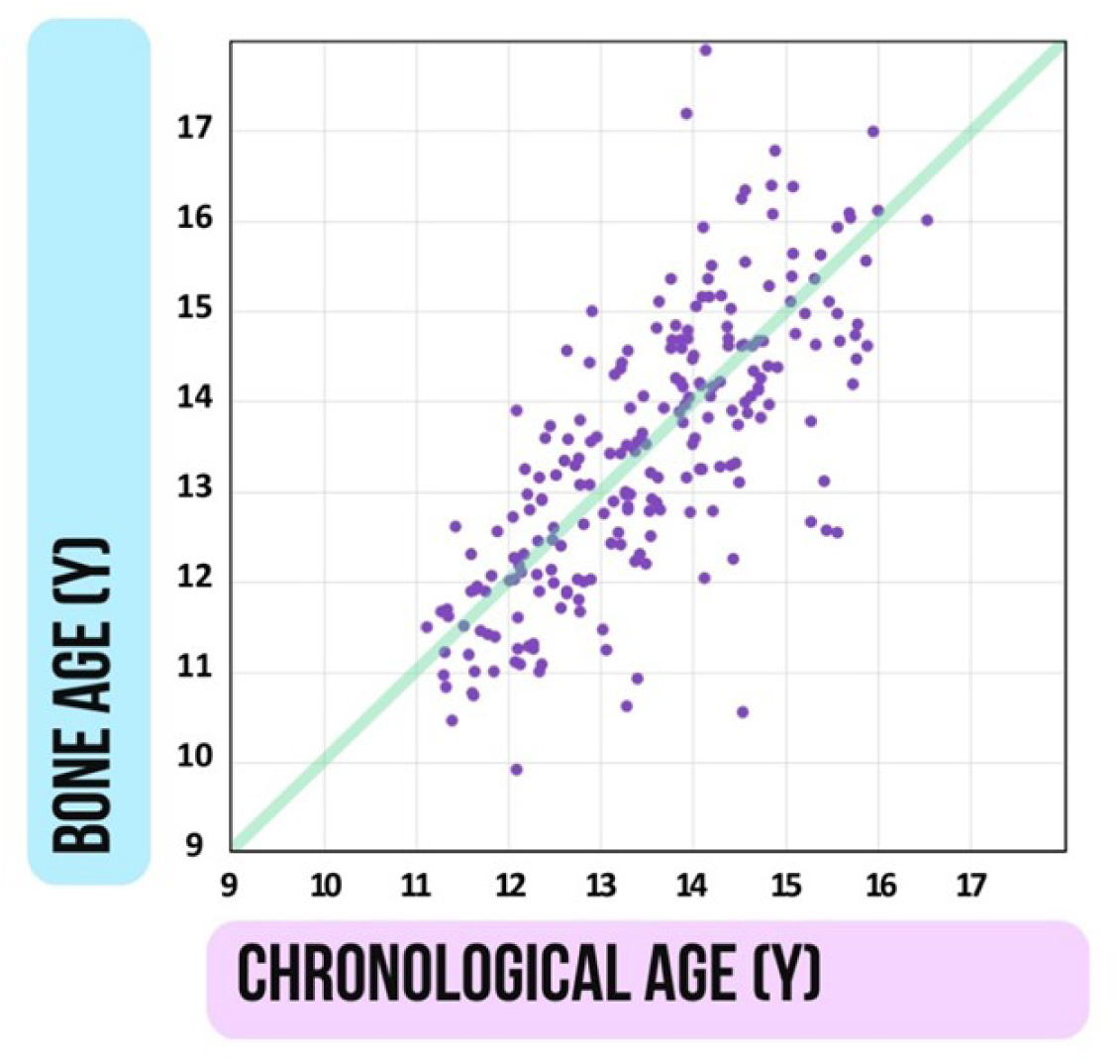
Variability of biological maturation relative to chronological age in the studied cohort. Biological maturity was assessed through bone age estimation (refer to Methods/Procedure). Both chronological age (CA) and bone age (BA) are expressed in years (Y), with each data point representing an individual in our sample. Along the diagonal green line, CA and BA coincide. Moving upwards away from the diagonal indicates accelerated maturation, while moving downwards indicates decelerated maturation. Although the two variables correlate significantly (r = .740, p < .001), the distribution of data points demonstrates that the disparity between CA and BA can exceed 2 years, signifying substantial differences in individual maturity levels (refer to Methods/Participants).

Lastly, while chronological age corresponds to the accumulation of experience, assessing individual experience levels is challenging. Fortunately, within the motor domain, a standardized practice regimen exists: instrumental musical practice. Considering the previously shown significant influence of musical experience on fine motor movement presented in Figure 2, Berencsi et al. ^35^, which could potentially obscure the effects of both biological maturity and chronological age, we analyze these effects within two distinct groups: individuals with musical experience and those without (see Figure 3.).

**Figure 3.**
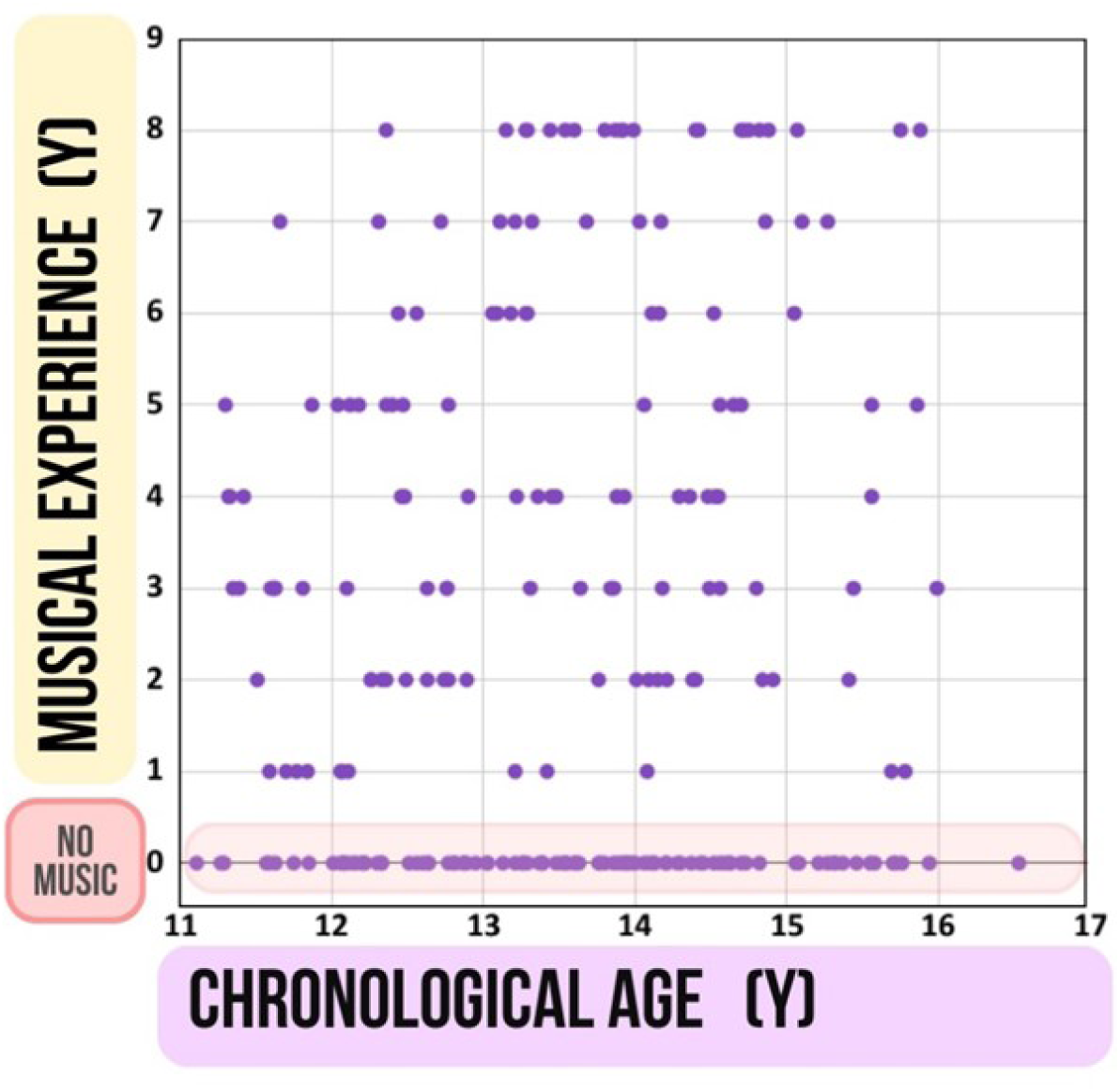
Two participant groups categorized based on instrumental musical experience, with one group comprising individuals who underwent specialized musical instrumental training (1-8 years). Musical experience, measured in years (Y), is plotted against chronological age in years (Y). Participants lacking instrumental musical experience are represented within the colored band at the bottom of the graph.

## RESULTS

In the subsequent two sections, we provide separate analyses for the participant groups distinguished by their musical experience status: those with musical experience and those without.

### Without musical experience

We conducted multiple linear regression analysis to predict performance on both sequential FT and index finger tapping tasks for both the dominant and non-dominant hand, utilizing BA and CA as predictors. The outcomes are outlined in Supplementary Table S3a. For the sequential FT task, a significant regression equation was found regarding the dominant hand (F(2,92) = 9.070, p < .001, R^2^ = .165) and also regarding the non-dominant hand (F(2,92) = 7.214, p < .001, R^2^ = .136). In both cases, only BA was a significant predictor of sequential finger tapping performance (p < .001). There were no significant regression equation regarding index FT (p > .05).

To avoid collinearity due to the high correlation of BA and CA, RWA was performed (Table 1.). In case of sequential FT performance, the relative weights of BA and CA were 79.76% and 20.24% on the dominant hand, and 73.29% and 26.71% on the non-dominant hand, respectively. The relative weights of BA were significant on both hands (p < .05). For index FT, relative weights of BA and CA were 37.55% and 62.45% on the dominant hand, and 16.34% and 83.66% on the non-dominant hand, respectively. The relative weights of CA were significant on both hands (p < .05). Results of RWA were in line with that of multiple regression analysis showing that BA is a significant predictor of sequential FT performance while index FT speed is predicted by CA in the group without musical experience.

**Table 1.**
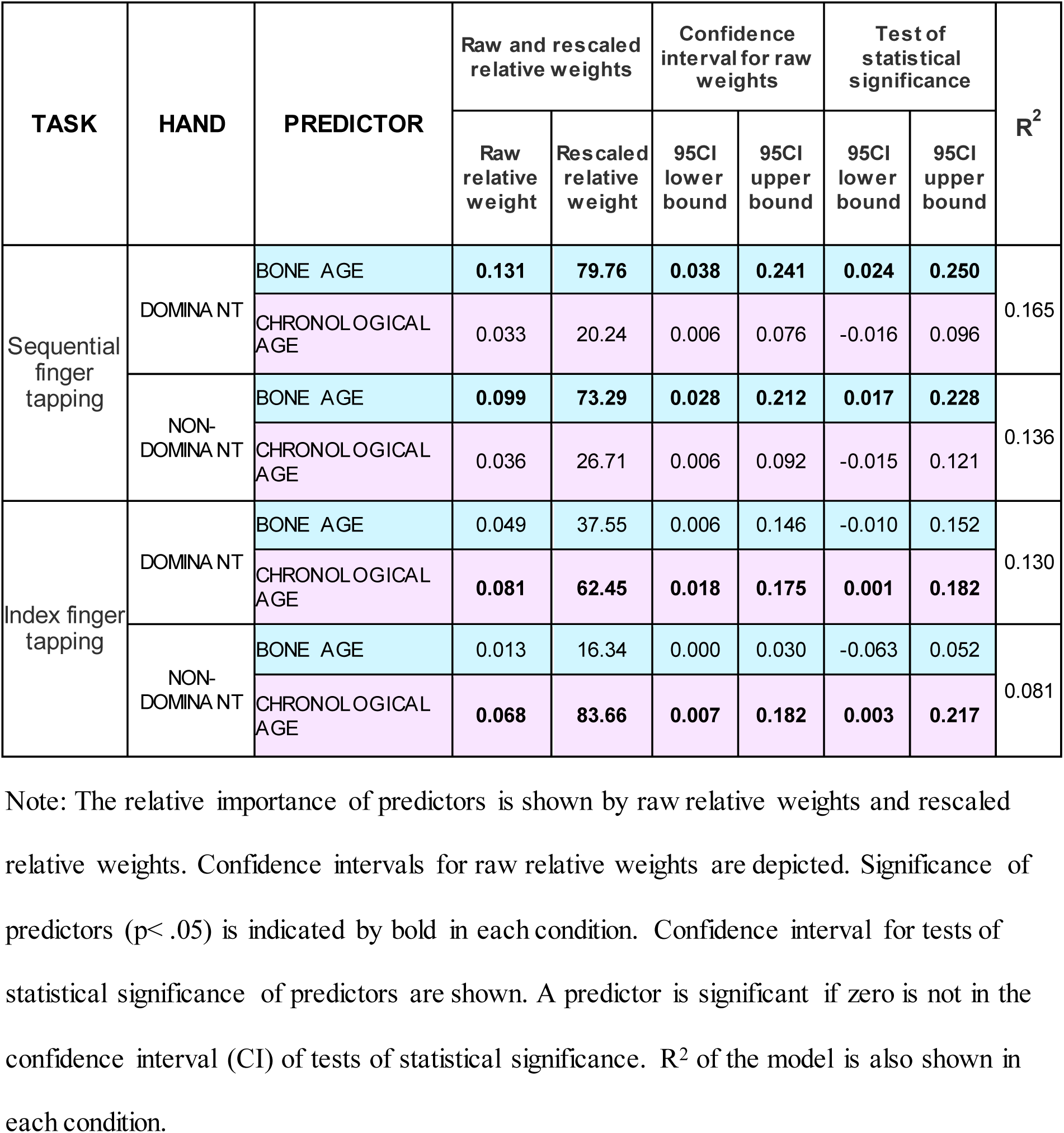
Results of RWA in the group without musical experience.

Predictor comparison shows if the relative weights from two predictors are significantly different from one another. Predictor comparison showed that BA was significantly different from CA as a predictor (p< .05) when performing the sequential FT task on the dominant hand. No other significant difference was present between predictors of CA and BA in the no musical experience group in sequential and index FT tasks (p> .05). Additional RWA analyses were performed to compare the relative weights between male and female participants. There was no significant difference between relative weights in either type of task by either hand (p> .05). Detailed results of the group without musical experience are shown in Supplementary Table S4a.

### With musical experience

We employed multiple linear regression analysis to predict performance in sequential and index FT tasks for both dominant and non-dominant hands, considering predictors including BA, CA, and instrumental musical experience. A comprehensive summary of the findings is provided in Supplementary Table S3b. We found a significant regression equation for the sequential FT task regarding the dominant hand (F(3,126) = 7.721, p < .001, R^2^ = .155) and the non-dominant hand (F(3,126) = 10.016, p < .001, R^2^ = .193). In both cases, the only significant predictor of sequential finger tapping performance (p ≤ .001) was instrumental musical experience. A significant regression equation was also found for the index FT task regarding the non-dominant hand (F(3,126) = 6.133, p = .001, R^2^ = .127). Instrumental musical experience was a significant predictor of non-dominant index FT performance (p < .05).

To avoid collinearity due to the high correlation of BA and CA, RWA was performed (Table 2). In case of sequential FT performance, RWA showed that relative weights of BA, CA and instrumental musical experience were 19.96%, 17.59% and 62.44% on the dominant hand, and 11.32%, 8.99% and 79.69% on the non-dominant hand, respectively. The relative weight of instrumental musical experience was significant on both hands (p < .05).

**Table 2.** Results of RWA in the group with musical experience.

| TASK | HAND | PREDICTOR | Raw and rescaled relative weights |  | Confidence interval for raw weights |  | Test of statistical significance |  | R<sup>2</sup> |
| --- | --- | --- | --- | --- | --- | --- | --- | --- | --- |
|  |  |  | Raw relative weight | Rescaled relative weight | 95CI lower bound | 95CI upper bound | 95CI lower bound | 95CI upper bound |  |
| Sequential finger tapping | DOMINANT | BONE AGE | 0.031 | 19.96 | 0.003 | 0.093 | -0.004 | 0.101 | 0.155 |
|  |  | CHRONOLOGICAL AGE | 0.027 | 17.59 | 0.004 | 0.087 | -0.004 | 0.106 |  |
|  |  | MUSICAL EXPERIENCE | **0.097** | **62.44** | **0.021** | **0.209** | **0.019** | **0.221** |  |
|  | NON-DOMINANT | BONE AGE | 0.022 | 11.32 | 0.002 | 0.073 | -0.019 | 0.079 | 0.193 |
|  |  | CHRONOLOGICAL AGE | 0.017 | 8.99 | 0.003 | 0.058 | -0.017 | 0.074 |  |
|  |  | MUSICAL EXPERIENCE | **0.153** | **79.69** | **0.053** | **0.279** | **0.055** | **0.289** |  |
| Index finger tapping | DOMINANT | BONE AGE | 0.024 | 59.55 | 0.002 | 0.091 | -0.071 | 0.101 | 0.040 |
|  |  | CHRONOLOGICAL AGE | 0.016 | 39.04 | 0.001 | 0.058 | -0.089 | 0.044 |  |
|  |  | MUSICAL EXPERIENCE | 0.001 | 1.41 | 0.000 | 0.001 | -0.102 | 0.017 |  |
|  | NON-DOMINANT | BONE AGE | 0.024 | 18.99 | 0.004 | 0.073 | -0.005 | 0.084 | 0.127 |
|  |  | CHRONOLOGICAL AGE | **0.048** | **37.62** | **0.004** | **0.122** | **0.002** | **0.145** |  |
|  |  | MUSICAL EXPERIENCE | **0.055** | **43.40** | **0.008** | **0.133** | **0.001** | **0.152** |  |

Note: The relative importance of predictors is shown by raw relative weights and rescaled relative weights. Confidence intervals for raw relative weights are depicted. Significance of predictors ( $p < .05$ ) is indicated by bold in each condition. Confidence interval for tests of statistical significance of predictors are shown. A predictor is significant if zero is not in the confidence interval (CI) of tests of statistical significance. R<sup>2</sup> of the model is also shown in each condition.

For index FT, relative weights of BA, CA and instrumental musical experience were 59.55%, 39.04 and 1.41% on the dominant hand, and 18.99%, 37.62% and 43.4% on the non-dominant hand, respectively. Relative weights did not reach significance with respect to the dominant hand, while relative weights of chronological age and instrumental musical experience were significant on the non-dominant hand (Table 2.). Results of RWA were in line with that of multiple regression analysis showing that instrumental musical experience is a significant predictor of sequential FT performance, while non-dominant hand index FT speed is predicted by chronological age and instrumental musical experience in the musical experience group.

Predictor comparison showed that musical instrumental experience differed significantly as a predictor from BA and CA during the execution of the sequential FT task with the non-dominant hand. No further comparison between the predictors of musical instrumental experience, BA and CA showed significant difference in sequential and index FT task (p> .05). Additional RWA analyses were performed to compare the relative weights between male and female participants. No significant difference between relative weights was found in either type of tasks by either hand (p> .05). Detailed results of the group with musical experience are shown in Supplementary Table S4b.

## DISCUSSION

In our exploration of the determinants of fine motor development in adolescence, the initial aim was to distinguish between the contributions of cortical network functioning and the degree of myelination in extended neural pathways. To achieve this, we utilized both a simple repetitive movement task and a sequential finger tapping task. To disentangle the potential influences of time (chronological age) and maturation (biological age), we utilized bone age as an indicator of individual maturation levels. Furthermore, to assess the impact of specialized experience and to prevent it from confounding the effects of chronological and biological age, we examined motor performance in two groups of adolescents: those with prior instrumental musical training and those without.

As summarized in the upper left panel of Figure 4., in the absence of musical experience, the main determinant of sequential finger tapping performance is bone age for both the dominant and the non-dominant hand. Conversely, for index finger tapping speed, chronological age emerges as the primary determinant for both hands. Notably, in the presence of musical experience, it becomes the predominant factor for sequential task performance on both hands. Regarding index finger tapping speed, musical experience influences performance solely on the non-dominant hand.

**Figure 4.**
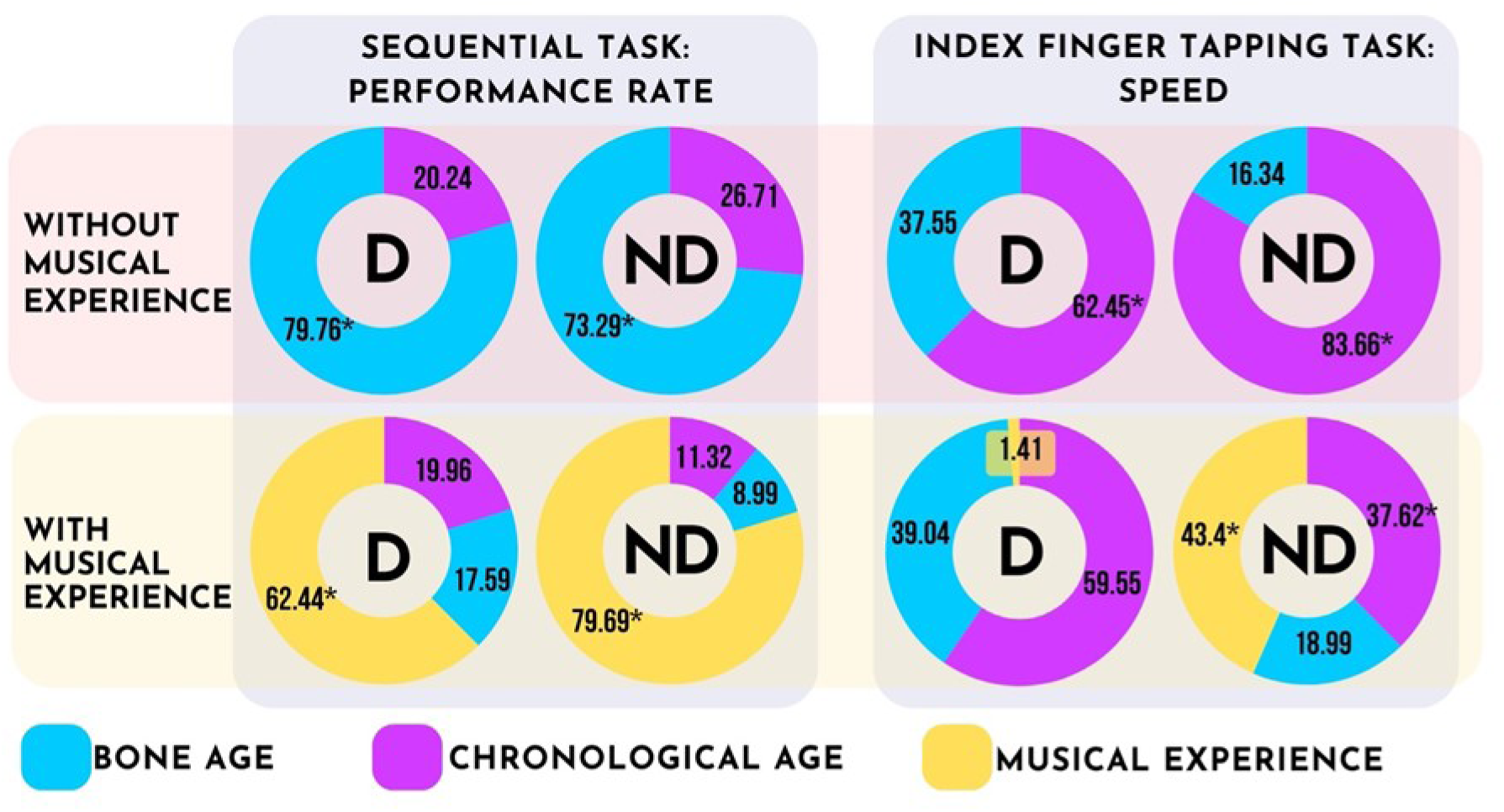
Graphical summary of results. D: dominant hand, ND: non-dominant hand. Charts show rescaled relative weights of predictors: bone age (blue), chronological age (magenta) and musical instrumental experience (yellow). Asterisks indicate significant predictors in the model (p < .05). When specific instrumental musical experience is not present, biological maturation level is a significant predictor of complex fine motor performance. On the other hand, simple repetitive motor performance is predicted by chronological age. With long-term instrumental musical motor experience, the amount of experience becomes the main predictor of complex fine motor performance and index finger tapping performance on the non-dominant hand.

Adolescent brain maturation involves intricate sequences of neurodevelopment ^10,42^, with normative trajectories indicating substantial volumetric changes ^43^ and synaptic density alterations ^44^ as depicted in Figure 5. These findings underscore the ongoing construction of the adolescent brain. One plausible interpretation of our primary results presented in the upper panels of Figure 4. suggests that while the speed of index finger tapping predominantly relies on the progressive enhancement of white matter myelination (illustrated by the gray curve in Figure 5), performance in sequential finger tapping tasks may be influenced by cortical regions undergoing maturation-dependent synaptic elimination processes (indicated by the purple curve in Figure 5).

**Figure 5.**
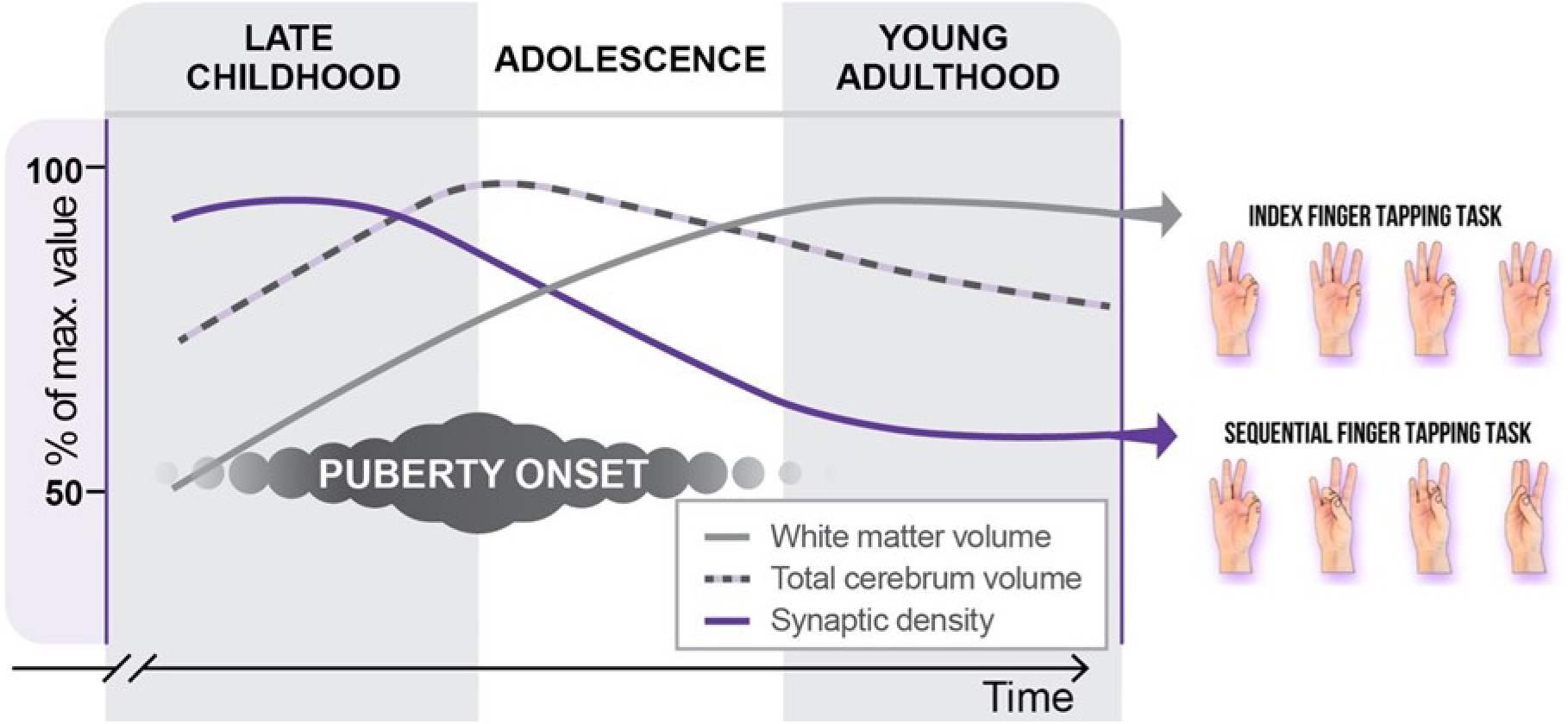
Interpretation of findings in the cohort without musical experience. The left panel illustrates schematic neurodevelopmental trajectories of the human brain. It is hypothesized that basic index finger tapping proficiency correlates with white matter myelination, a process dependent on time, thereby influencing the speed of neural transmission. On the other hand, performance in sequential finger tapping tasks is postulated to be influenced by maturation-dependent synaptic elimination, facilitating more efficient network interactions within the motor cortices.

We have shown that chronological age is the main determinant of repetitive fine motor speed when musical experience is not present. This result might be related to the long-term, chronological age associated processes of white matter development including myelination and axon caliber increase of the descending tracts ^7,16^, and white matter development in motor related cortical pathways ^45^. The latter parallels the U-shaped distribution of repetitive fine motor speed as a function of chronological age with a peak at around 38 years ^13,45,46^. On the other hand, structural variations of the corticospinal tract related to sex hormone levels are not necessarily reflected in adolescent motor performance ^40^, however these structural changes may precede and establish the basis for experience-dependent development of behavioral performance ^15^.

Bone age associated sequential fine motor performance likely reflects pubertal maturity related cortical processes. Trajectories of puberty related cortical thinning in frontal and parietal cortices show association with pubertal tempo, suggesting an independent process from merely age-related development ^18^. Targeted elimination of synapses or synaptic pruning ^44,47–49^ that occurs throughout the nervous system during development also seems to be linked to puberty onset in rodents ^23,50^ and humans ^44^. In the human primary motor cortex, synaptic density decreases until late adolescence ^9^ indicating a prolonged formation of efficient cortical networks by pruning ^51^. Here we showed the influence of biological maturation on the complex fine motor performance during adolescence for the first time and hypothesize that it might be linked to the mentioned puberty related cortical developmental events.

In the presence of highly specific motor experience, the effect of biological maturation diminishes, and the amount of experience becomes the main predictor of complex fine motor performance (Figure 4.). In a similar study that did not make a distinction between participant groups with or without musical instrumental experience (that is, the analyzed cohort included both groups of participants) the effect of musical instrumental experience was also superior to bone age and chronological age ^35^. This is explained by earlier studies showing that long-term motor experience shapes motor cortical areas both in terms of structural changes ^52^ and at the functional network level ^38,53,54^, paralleling behavioral improvements in fine motor performance ^8,38,55,56^. Fine motor experience may shape motor cortical areas through synaptic pruning, leading to more efficient cortical networks ^57^. The prolonged development of long-range cortico-cortical connections that are fundamental to complex motor performance ^58^ are also prone to experience-dependent alterations ^52,59^. When comparing the dominant and the non-dominant hands, determinants of hand function are largely consistent between the two body sides (Supplement S4) despite the superior performance of the dominant hand in both motor tasks (Supplement S5). In the absence of musical instrumental experience, the main determinant of sequential motor performance was bone age but musical instrumental experience shaped the determinants of complex fine motor performance very comparable on both sides (Figure 4.) in a laterality-independent way.

In this study we successfully dissociated the effect of biological maturation on the determinants of fine motor performance, however, applying inclusion criteria regarding age, education, and socioeconomic status may pose limitations with respect to the generalizability of our results. On the other hand, these restrictions were necessary to avoid confounders in a cross-sectional study with a focus on maturation and experience. At the same time, inclusion of a considerable number (n=255) of both male and female participants allows for generalizability to each sex.

Further exploring the effect of cultural background and environmental factors such as device use in terms of touch screens or game controllers may add to the understanding of hand function in modern society.

In the current study we focused on adolescence, however, opening up the investigated developmental window towards childhood in future longitudinal assessments may also reveal the interplay between biological maturation and early experience during the development of neural networks and hand function. Furthermore, populations with neurodevelopmental disorders often show impaired fine motor skills ^37,60^ and may also show differential pubertal maturation patterns, e.g., precocious puberty ^61,62^. The mapping of association between brain functional alterations and fine motor learning capacity in atypical development is rare ^63^, and its relation to pubertal development still needs to be addressed.

## CONCLUSIONS

In conclusion, by disentangling the effects of biological maturation and experience during adolescence, we found that biological maturation is a significant contributor to complex fine motor performance when highly specific instrumental musical experience is not present. In terms of the neural background, maturation-dependent synaptic elimination processes of the related cerebral cortical regions are suspected. In the case of simple repetitive fine motor behavior, we found that chronological age is the main contributor, which suggests the influence of age-dependent maturation of white matter. In the presence of highly specific musical instrumental experience, the amount of experience becomes the main contributor to both complex sequential and simple repetitive performance. Our findings emphasize both the important role that biological maturation plays in the development of complex fine motor behavior, and also the relevance of experience dependent behavioral plasticity.

## METHODS

### Participants

Participants were recruited by contacting schools in high-level socioeconomic regions in Budapest, Hungary, as well as through online advertisements. Parents provided demographic, educational, and medical history details. Exclusion criteria included the presence of learning disability, sleep disorder, developmental disorder, neurological disorder, or injury affecting the function of upper extremities, based on a questionnaire filled by parents. Initial enrollment was 269 participants without the above-mentioned exclusions. We then excluded 44 further participants due to the lack of reporting musical experience (N = 15), performance on the motor task not reaching 50% accuracy (N = 13), not completing all tasks or equipment malfunction (N =14), reporting mixed handedness (N = 1), and reporting ADHD diagnosis after enrollment (N =1). After exclusions, 225 adolescent participants took part in the motor development study (F = 123). Chronological age (CA) ranged between 11.1 and 16.5 years (M = 13.5, SD = 1.2) and bone age ranged between 9.9 and 17.9 years (M = 13.5, SD = 1.5).

The group without instrumental musical experience was comprised of 95 participants (F = 39, 11.1-16.6, M = 13.6, SD = 1.2). The group with instrumental musical experience was comprised of 130 participants (F = 84, CA range 11.3-16.0, M = 13.4, SD = 1.2). 28 of the participants were left-handed based on the reports of hand preferred to perform unimanual tasks. For a detailed description of the participant groups, see Supplement S1.

Motor development data collection was part of a large-scale study, where each participant was also administered the Wechsler Intelligence Scale for Children, Fourth Edition ^24^, carried out a Stroop test ^64^, and provided resting state EEG recordings ^36^.

The Hungarian United Ethical Review Committee for Research in Psychology (EPKEB) approved the study (reference number 2017/84). Written informed consent was obtained from all the subjects and their parents. Participants were given gift cards (approx. EUR 15 value each) for their attendance. All research described in this paper was performed in accordance with the Declaration of Helsinki.

### Procedure

#### Bone age assessment

Bone age measurement was used to determine the biological development of participants as described in earlier studies ^24,26,35,36^. This method measures the process of ossification in the wrist and hand to estimate the bone age of the individual. A standard anthropometer (DKSH Switzerland Ltd, Zurich, Switzerland) was employed to measure body height, and body mass was measured using a scale (Seca). An ultrasound-based device (Sunlight Medical Ltd, Tel Aviv, Israel) was used to estimate bone age. Using this measurement, the ultrasound passes through the subject’s left wrist, evaluating the distal epiphyseal and diaphyseal ossification of the two forearm bones (radius and ulna). The device measures the speed of ultrasound and the distance between the transducers (wrist width) to estimate bone age (in years and months).

Measurements were repeated five times with the transducers being raised 2 mm for each measurement. Bone age data were corrected using a standard Hungarian bone age database (for details see Supplement 2). Using ultrasound for bone age assessment is safe for the subjects as opposed to ionizing radiation-based techniques, while results obtained from these two types of procedures are highly correlated ^65^.

#### Assessment of instrumental musical experience

A questionnaire was administered to gather data on participants’ musical instrumental experience. Participants were asked to indicate the number of years spent learning an instrument, with the total years of training across multiple instruments computed if applicable. The cumulative years of playing one or multiple instruments ranged from 0 to 8 years (Experience 0 - 8 years, M = 13.5, SD = 1.5). There were no restrictions regarding the type of musical instrument. In Hungary, musical instrumental education is broadly accessible in the form of music schools. Music schools have a standard schedule of afternoon lessons of 45-minute solfeggio and 30-minute instrumental lessons twice a week ^66^.

Participants were categorized into two groups based on their musical instrumental experience (Figure 3.). The group without musical experience (N = 95) comprised of individuals with either no musical experience or less than one year of experience. The group with musical experience (N = 130) consisted of participants with at least one year and up to a maximum of eight years of musical instrumental experience (see Figure 3.).

#### Assessment of fine motor development

Motor tasks depicted in Figure 1. and previously used in developmental studies ^3,35,37,63^ were executed using both the dominant and non-dominant hands. Initially, all participants undertook the task with their non-dominant hands. The sequential finger tapping (FT) task entailed a four-element finger-to-thumb opposition sequence involving the index, ring, middle, and little fingers. Each participant completed ten blocks of 16 repetitions, aiming for maximal speed and accuracy. Data collection commenced once participants successfully executed three consecutive sequences accurately with closed eyes. To effectively balance the speed-accuracy trade-off inherent in this task, the dependent variable was defined as Performance Rate (PR), calculated as the number of correctly performed finger taps per second (taps/s). The mean PR across ten blocks for each participant was utilized in statistical analyses.

The non-sequential task involved an index finger to thumb opposition maneuver. Participants executed 64 taps at maximum speed. The dependent variable was defined as the number of taps per second (taps/s). The mean number of taps across three blocks of 64 taps was computed for each participant.

A custom-designed data glove was utilized to capture the sequence and timing of finger tapping. This glove secured electrodes on the palmar surface of the distal phalanges of each finger. Connected via a serial-to-USB hub, the data glove interfaced with a laptop computer. A custom Java-based software was employed to collect data on the timing and sequence of finger taps, as well as to automatically compute the Performance Rate (PR) and number of index finger taps per second for each participant.

#### Statistical analysis

The sample was stratified into two groups based on instrumental experience: a group without musical experience and a group with musical experience. Subsequent analyses were conducted separately for each group. This approach is supported by earlier findings showing that musical instrumental experience may be an extremely strong determinant of fine motor performance, potentially masking the effect of bone age ^35^.

A two-step analysis was executed for each dependent variable (performance rate and maximum motor speed) for both the dominant and non-dominant hand. First, a multiple regression analysis was carried out to elucidate the independent effects of chronological age (CA) and bone age (BA) in both groups. In the group with musical experience, the duration of musical instrumental experience was also incorporated into the regression model. Thus, four models were constructed for each group (two models for each finger tapping variable of the dominant and non-dominant hands), yielding a total of eight models. Given the high positive correlation between biological age and chronological age (r (225) = .740, p < .001), collinearity issues arose, necessitating further analysis.

Relative weight analysis (RWA) was employed to determine whether the predictors tapped into the same variance. Four separate RWAs were conducted for each group (two for each finger tapping variable of the dominant and non-dominant hands) to assess whether the weights of predictors significantly differed from one another. Additionally, supplementary RWAs were performed (1) to examine whether relative weights differed from each other in each condition (FT task×hand) by pairwise comparisons between predictors in the model and (2) to examine whether weights differed significantly between male and female groups (two analyses for each finger tapping variable of the dominant and non-dominant hands in each group).

Multiple regression analysis was performed by IBM SPSS Statistics for Windows, Version 23.0. (IBM, Armonk, New York), and RWA was performed by WA-Web App ^67^. The significance level was set at p< .05.

## ACKNOWLEDGEMENTS

We thank the generous and amazing parents, adolescents and schools who participated in this project.

The project was supported by the National Research, Development and Innovation Office of Hungary (Grant K-134370 to I.K.) and by the Hungarian Research Network (HUN-REN-ELTE-PPKE Adolescent Development Research Group). It was made in the framework of the PPKE-BTK-KUT-23-1 project, with the support and funding provided by the Faculty of Humanities and Social Sciences of Pázmány Péter Catholic University. The funders had no role in study design, data collection and analysis, decision to publish or preparation of the manuscript.

## AUTHOR CONTRIBUTIONS

I.K., B.A., F.G. and P.G. designed the experiment; G.O. recruited and screened participants, K.U. and Z.T. collected and analyzed bone age data, B.A. analyzed the finger tapping data; B.A., K.I. and L.J.F. drafted the paper and all authors contributed to discussion and to the text. All authors approved the final version of the manuscript.

## DATA AVAILABILITY

The data necessary to reproduce the analyses presented here are not publicly accessible but available from the corresponding author upon reasonable request. The analytic code necessary to reproduce the analyses presented in this paper is publicly accessible. The analyses presented here were not preregistered.

## SUPPLEMENTARY MATERIAL

Participant descriptives and further analyses are available at the Open Science Framework (OSF) platform at this address: https://osf.io/zyvxn/?view_only=74c6c9233d6343f5aee207cffe2c0868

## ADDITIONAL INFORMATION

The authors have no conflict of interest to disclose.

## Abbreviations

BA: bone age
CA: chronological age
FT: finger tapping

## Notes

https://osf.io/zyvxn/?view_only=74c6c9233d6343f5aee207cffe2c0868

### Competing Interest Statement

The authors have declared no competing interest.

